# Molecular basis of stepwise cyclic tetra-adenylate cleavage by the type III CRISPR ring nuclease Crn1/Sso2081

**DOI:** 10.1101/2022.10.01.510428

**Authors:** Liyang Du, Zhipu Luo, Zhonghui Lin

## Abstract

The cyclic oligoadenylates (cOAs) act as second messengers of type III CRISPR immunity system through activating the auxiliary nucleases for indiscriminate RNA degradation. The cOA-degrading nucleases (ring nucleases) provide an ‘off-switch’ regulation of the signaling, thereby preventing cell dormancy or cell death. Here, we describe the crystal structures of the CRISPR-associated ring nuclease 1 (Crn1) from *Saccharolobus solfataricus* (Sso) 2081 in its apo or bound to cA_4_ in both pre-cleavage and transient intermediate states. Sso2081 harbors a unique helical insert that encloses cA_4_ in the central cavity. Two free phosphates symmetrically bind the catalytic site of apo Sso2081 and overlap with the two scissile phosphates of cA_4_, supporting a bilaterally symmetrical cleavage. The structure of transient intermediate state captured by Ser11Ala mutation immediately illustrates a stepwise cleavage of cA_4_ by Sso2081. Our study establishes atomic mechanisms of cA_4_ recognition and degradation by the type III CRISPR ring nuclease Crn1/Sso2081.

## INTRODUCTION

The CRISPR system provides adaptive immunity against mobile genetic elements in bacteria and archaea (1-3). Upon the invasion of foreign DNA and / or RNA, the surveillance systems are activated to degrade the invaders through the ribonucleoprotein (RNP) complexes (4-7). According to the composition of RNP complexes, CRISPR systems can be divided into two classes (8,9). The class 1 systems (types I, III, and IV) consist of multi-subunit RNPs (10), whereas the class 2 systems (types II, V, and VI) possess single-subunit RNPs (11,12).

The Type III CRISPR systems are featured by the presence of Cas10 signature protein (also named as Csm1 or Cmr2), which contains an HD nuclease domain for ssDNA cleavage and a cyclase domain for cyclic oligoadenylate (cOA) synthesis (13-17). The recognition of a target RNA by RNP complex stimulates the cyclase domain to synthesize cOA molecules. These cOA molecules (typically ranging 3 to 6 AMPs), as second messengers, in turn allosterically activate the CRISPR ancillary nucleases such as Csx1/Csm6 (18,19), Can1/Can2 (CRISPR ancillary nuclease 1/2) (20,21), and Card1 (cyclic-oligoadenylate-activated single-stranded ribonuclease and single-stranded deoxyribonuclease 1) (22), resulting in indiscriminate degradation of both foreign and host DNA and / or RNA(23,24). Therefore, although cOA is critical for host immunity against foreign genetic elements, its cellular level must be tightly controlled so as to avoid cell dormancy or cell death (25).

Recently, a group of CRISPR-associated Rossman-fold (CARF) domain containing proteins, termed the ring nucleases, have been described to cleave cOA molecules in a metal-independent mechanism (26-32). Based on the functionality, these ring nucleases can be divided into two major categories: (i) the standalone ring nucleases that are specific for cOA degradation, such as the *S. solfataricus* (Sso) Sso2081 (26) and *S. islandicus* (Sis) Sis0811 (Crn1) (27), the anti-CRISPR (Acr) III-1 (Crn2) (28), and Csx3 (Crn3) (29); (ii) the self-limiting cOA-dependent ribonucleases like Csm6 from *T. onnurineus* (30), *T. thermophilus* (31) *and E. italicus* (32), which also consist of a HEPN (higher eukaryotes and prokaryotes nucleotide) domain for DNA and / or RNA degradation.

Sso2081 is the first member of ring nuclease family identified from *S. solfataricus* (26). It has been shown that Sso2081 could convert cA_4_ into 5□-OH-ApA-2□, 3□-cyclic phosphate (A_2_>P), and hence inactivate the RNase activity of Csx1 in target RNA clearance (26). These findings represent an important milestone in our understanding of the regulations of CRISPR system, however, the structural mechanisms of cA_4_ recognition and cleavage by Sso2081 remain to be established. In the present work, we have determined the crystal structures of Sso2081, alone or bound to cA_4_ in both pre-cleavage and transient intermediate states. These structures together with extensive biochemical analyses extend our understanding of the molecular basis of ‘off-switch’ regulation for the CRISPR system.

## MATERIALS AND METHODS

### Oligonucleotides and cloning

cA_4_ (Cyclic tetraadenosine monophosphate) was ordered from Biolog Life Science Institute, Bremen, Germany. The Sso2081 cDNA (GenBank ID: 1559988754) was synthesized at GenScript Corporation (Nanjing, China). The cDNA of Sso2081 and its variants were subcloned into a modified pET bacterial expression vector with an N-terminal cleavable His_6_-tag.

### Protein expression and purification

The Sso2081 protein was expressed in *Escherichia coli* Rosetta (DE3) cells. Cells cultured to OD_600_ ∼ 0.6 were induced with 0.5 mM isopropyl β-D-1- thiogalactopyranoside (IPTG) at 18□ overnight. Then, the cells were harvested, resuspended with lysis buffer (20 mM Tris-HCl pH 8.0, 200 mM NaCl, 10 mM imidazole, 5% glycerol, and 1‰ Tween-20), and disrupted by French Pressure (Union Biotech, China). The His_6_-tagged protein in the supernatant was pooled through Ni-NTA resin (Union Biotech, China). The column was subjected to extensive wash with 40 mM imidazole containing lysis buffer, and the protein of Sso2081 was eluted with lysis buffer supplemented with 200 mM imidazole. After removal of His_6_ tag with the home-made preScission protease, the untagged protein was further purified through 15Q anion exchange column (GE Healthcare Life Sciences) and Uniondex 200 pg 16/60 size-exclusion column (Union Biotech, China). The final purified protein in 20 mM Tris-HCl pH 8.0 and 150 mM NaCl was concentrated and stored at −80□.

The selenomethionine (SeMet) labelled protein of Sso2081 was produced as previously described (33). Briefly, the Rosetta (DE3) cells containing pET-Sso2081 plasmid were cultured in Luria-Bertani (LB) media overnight. The cells were collected and washed with M9 minimal media, and further cultured in M9 minimal media at 37 °C until the OD_600_ reached about 0.6. Then, the amino acid mixture containing 50 mg/L of leucine, isoleucine and valine, 100 mg/L of phenylalanine, lysine and threonine, and 80 mg/L of SeMet was added to the culture. Protein expression was then induced with 0.5 mM IPTG at 18 °C overnight. The SeMet labelled protein was subsequently purified using the same protocol as described above.

### cA_4_ cleavage assay

A cA_4_ cleavage assay was conducted to determine the ring nuclease activities of Sso2081 and its variants. In a 50-μL reaction, 40 μM of synthetic cA_4_ was incubated with 4 μM or indicated concentrations of Sso2081 in the cleavage buffer containing 20 mM Tris-HCl pH 8.0 and 50 mM NaCl at 37 °C for 15 min or indicated time. At the end of the reaction, 50 μL chloroform-isoamylol (24:1) was added, followed by vortexing for 60 s. Then the samples were centrifuged at 10,000 rpm for 8min. After another round of chloroform-isoamylol extraction, the top aqueous phase containing nucleotides were collected for further analyses on liquid chromatography (LC) and mass spectrometry (MS).

### LC-MS analyses

The cA_4_ and its cleavage products extracted by chloroform-isoamylol were separated using a High Performance Liquid Chromatography (HPLC) system (LC-20A, Shimadzu) equipped with a RX - C18 column (2.1 × 100 mm, 5 μm,Zhongpu Science). After injection of 20 μL sample, the column was eluted with a linear gradient of buffer-B (acetonitrile supplemented with 0.01% TFA) against buffer-A (water with 0.01% TFA) at a flow rate of 0.35 mL / min as follows: 0 - 2 min, 2 - 20% B; 2 - 5 min, 20% B; 5 - 12 min, 20%-48% B; 12 - 13 min 48–95% B; 13 - 25 min 95% B; 25 – 26 min, 95% - 2% B; 26 – 35 min, 2% B. The column temperature was set to 40 □ and the UV data were recorded with a wavelength of 259 nm. Mass spectra data were acquired in negative-ion mode with scan range of *m/z* 150 - 1500 on an Agilent 6520 ACURATE-Mass Q - TOF mass spectrometer. The mass spectrometer was operated in full scan and multiple reaction monitoring (MRM) modes. The capillary voltage was 3.5 kV and the temperature was 350 [. Nebulizer pressure was set to 40 psi, and the drying gas flow rate was 10 L/min. Nitrogen was used as nebulizer and auxiliary gas.

### Crystallization and data collection

Crystallizations were performed at 25 °C using the hanging-drop vapor diffusion method. Crystallization drops were set up by mixing protein or protein-ligand complex with equal volume of reservoir solutions. Crystals were cryoprotected by the reservoir solutions supplemented with 15%-25% glycerol prior to data collection.

The crystals of SeMet-Sso2081 were grown with 20 mg/mL of protein in the reservoir solution consisting of 0.1 M sodium acetate trihydrate pH 4.5 and 25% w/v polyethylene glycol 1500. For the crystallization of Sso2081:cA_4_ complex, 16 mg/mL of Sso2081 protein was pre-incubated with cA_4_ at 4 °C with a molar ratio of 1:1.5, the crystals were grown with a reservoir solution containing 0.1 M sodium acetate trihydrate pH 5.0, 22% w/v polyethylene glycol monomethyl ether 550 and 5% w/v n-dodecyl-β-D-maltoside. For the crystallization of Sso2081 Ser11Ala : A_4_>P complex, 16 mg/mL of Sso2081^S11A^ protein was pre-incubated with cA_4_ at 25 °C with a molar ratio of 1:1.5, crystals were obtained using the reservoir solution containing 0.1 M sodium acetate trihydrate, pH 5.0 and 20% w/v polyethylene glycol 1500.

X-ray diffraction data of the crystals of SeMet-Sso2081 and Sso2081 Ser11Ala : A_4_>P were collected at beamline of BL02U1 with a wavelength of 0.979 Å, and the data of Sso2081:cA_4_ were collected at BL19U1 with a wavelength of 0.978 Å, at National Facility for Protein Science in Shanghai (NFPS), at Shanghai Synchrotron Radiation Facility (SSRF). Diffracting data were processed with XDS (34) and HKL2000 software (35).

### Structure determination and refinement

The initial phase for structure determination was obtained by the selenium single anomalous dispersion (SAD) method. The crystal of SeMet-Sso2081 diffracted to 2.70 Å resolution and exhibited the symmetry of the space group *P*2_1_ with cell dimensions of a = 67.03 Å, b = 38.38 Å and c = 69.80 Å; α = 90.00º, β = 106.41º and γ = 90.00º. Using the data truncated to 2.70 Å, 8 possible selenium sites were located and refined with the Autosol program in the PHENIX package (36), resulting in an overall figure of merit of 0.31. The resulting electron density map was used to construct an initial model with the AutoBuild program in PHENIX (37).

The structures of Sso2081:cA_4_ and Sso2081^S11A^: A_4_>P complexes were solved by molecular replacement with the program Phaser-MR of PHENIX (38), using the structure of apo Sso2081 as template. The resulting solution was used to construct the initial model with the program AutoBuild in PHENIX (37). Iterative model building and refinement were carried out with COOT (39), PHENIX (36) and REFMAC5 (40). The final models were validated with the program MolProbity in PHENIX (36). Data collection and refinement statistics were summarized in Table. 1. All structural figures in this study were generated with the PyMOL program (http://www.pymol.org/).

**Table 1.** Data collection and structure refinement statistics.

|  | SeMet-Sso2081 | Sso2081:cA<sub>4</sub> | Sso2081<br>S11A:cA<sub>4</sub>>P |
| --- | --- | --- | --- |
| **PDB ID** | **7YHL** | **7YGH** | **7YGL** |
| **Data Collection** |  |  |  |
| Space group | *P*2<sub>1</sub> | *P*2<sub>1</sub> | *P*2<sub>1</sub> |
| Cell dimensions |  |  |  |
| *a*, *b*, *c* (Å) | 67.03, 38.38, 69.80 | 67.05, 38.84, 72.97 | 67.51, 39.14, 74.09 |
| $\alpha$, $\beta$, $\gamma$ (°) | 90, 106.4, 90 | 90, 105.8, 90 | 90, 105.2, 90 |
| Resolution (Å) | 40.95-2.70<br>(2.80-2.70)* | 50.00-3.11<br>(3.22-3.11) | 50.00-2.50<br>(2.59-2.50) |
| *R*<sub>merge</sub> (%) | 6.7 (74.8) | 7.0 (78.6) | 4.3 (85.1) |
| (*I*)/$\sigma$(*I*) | 10.9 (1.5) | 22.84 (1.89) | 32.22 (1.86) |
| Completeness (%) | 93.9 (95.6) | 99.57 (97.89) | 99.9 (92.6) |
| Redundancy | 5.3 (5.3) | 3.7 (3.5) | 6.4 (5.8) |
| **Refinement** |  |  |  |
| Resolution (Å) | 40.95-2.70 | 28.56-3.11 | 28.91-2.50 |
| No. reflections | 8615 | 6382 | 12407 |
| *R*<sub>work</sub> / *R*<sub>free</sub> | 0.263/0.288 | 0.238/0.258 | 0.258/0.279 |
| No. atoms | 2710 | 2937 | 2956 |
| Macromolecules | 2641 | 2816 | 2822 |
| Ligand/ion | 66 | 88 | 88 |
| Water | 3 | 1 | 6 |
| *B*-factors | 124.3 | 139.9 | 117.7 |
| Macromolecules | 124.3 | 140.9 | 118.6 |
| Ligand/ion | 126.7 | 109.4 | 90.7 |
| Water | 67.3 | 42.7 | 83.4 |
| R.m.s. deviations |  |  |  |
| Bond lengths (Å) | 0.010 | 0.010 | 0.010 |
| Bond angles (°) | 1.80 | 1.71 | 1.76 |
| Ramachandran plot (%) |  |  |  |
| Favored/allowed/disallowed | 94.41/5.59/0.00 | 93.58/6.42/0.00 | 96.09/3.91/0.00 |

\*Values in parentheses are for highest-resolution shell.

## RESULTS

### Crystal structure of Sso2081

Sso2081 is a standalone ring nuclease with single CARF domain and functions as a homodimer. For crystallization, we expressed and purified the full length protein (aa.1-178) from bacterial cells. We first tested whether the recombinant protein of Sso2081 was active in cA_4_ cleavage by high-performance liquid chromatography-mass spectrometry (HPLC-MS) analyses. The recombinant Sso2081 could cleave synthetic cA_4_ in both time- and concentration- dependent manners (Figures 1A and 1B, Supplementary Fig. 1). Mass spectrometry analysis showed that Sso2081 converted the cyclic cA_4_ into linear A_2_>P (Supplementary Fig. 1), yielding a single-turnover rate constant of ∼ 0.83 min^-1^, which is comparable to the previous results (0.23 min^-1^) (26).

**Figure 1.**
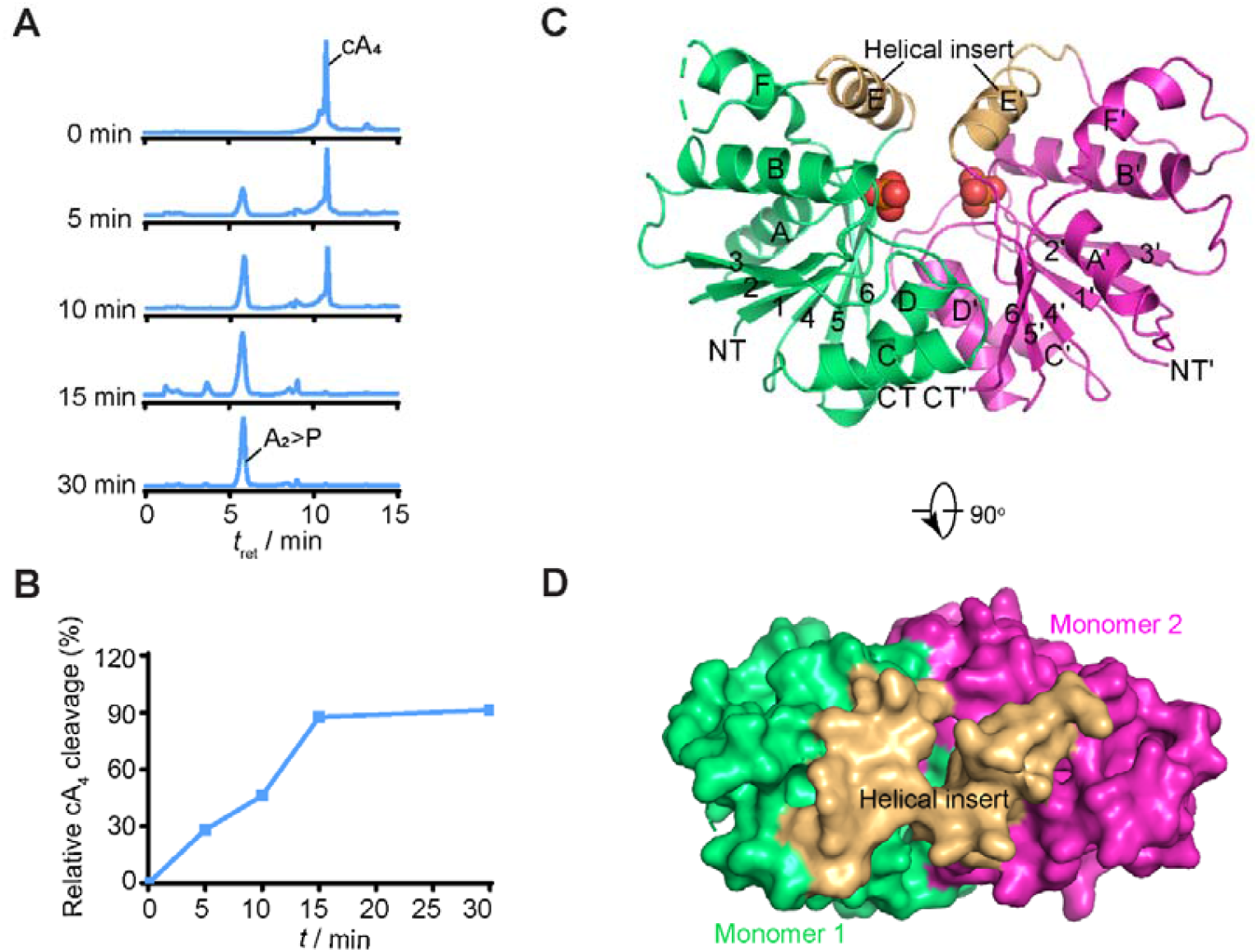
Crystal structure of the apo Sso2081. **(A)** LC-MS based real-time monitoring of cA_4_ cleavage by Sso2081. The mass spectra of cA_4_ and cleaved products are shown in Supplementary Fig. 1. **(B)** The kinetic plot of cA_4_ cleavage in (A). Values are means of duplicate measurements. **(C)** The overall structure of Sso2081. The structure is shown in cartoon representation and the bound free phosphates are in spheres. The two homo-monomers are colored in green and magenta, and helical inserts are highlighted in yellow. The secondary structural elements are labeled. The N and C termini are indicated as NT and CT. **(D)** Top-view of the Sso2081 structure overlaid with a 60% transparent surface. All structural figures in this study, unless otherwise indicated, use the same color and labeling schemes.

We next crystallized the native protein of Sso2081. However, presumably owing to limited sequence identity to its structural homologs (Supplementary Fig. 2), we were unable to determine the structure by molecular replacement. We then tried to determine the structure by single-wavelength anomalous dispersion (SAD) method. The selenomethionine (SeMet)-labeled Sso2081 crystals diffracted to a minimum Bragg spacing of 2.70 Å resolution. The structure was solved using the Autosol program in the PHENIX package (36). The final refined model contains two molecules in the asymmetric unit with good geometry. Data collection and refinement parameters were summarized in Table. 1.

The structure of apo Sso2081 reveals a butterfly-shaped homodimer in a two-fold symmetry (Figure 1C). In each monomer, there are six β-strands sandwiched by six α-helices with four on one side and two on the other side, forming a canonical Rossman fold. The dimerization of the two monomers mainly involves helices C, D and β6, which together generate a buried surface of about 1198.7 Å^2^. A structural homology search with the DALI server (41) revealed that Sso2081 is structurally related to the CARF domains of Can2 (PDB: 7BDV, RMSD = 2.2 Å) (21), Card1 (PDB: 6WXX, RMSD = 2.6 Å) (22), and TtCsm6 (PDB: 5FSH, RMSD = 2.4 Å) (18) (Supplementary Fig. 3). Despite the similarity, Sso2081 also displays some unique structural features. For example, it possesses six β-strands, whereas others have seven (Supplementary Fig. 3). The most prominent feature of Sso2081 is the presence of a helical insert (αE) between β5 and β6 (residues 130-168), which hangs over the dimer interface and shields the catalytic center (Fig. 1c and 1d). On the contrary, the active sites within the CARF domains of Can2, Card1 and Csm6 are generally in an open conformation (Supplementary Fig. 3).

### The free phosphate-binding sites

Like many other ring nucleases, the catalytic pocket of Sso2081 is located above the dimer interface, featured by a central positively charged region (Figure 2A). This basic patch likely binds the phosphodiester backbone of cA_4_ substrates as shown in the previous structures of cA_4_-bound CARF domains. Notably, two free phosphates, presumably from the protein purification process, bind two symmetrical catalytic pockets with unambiguous electron density (Figures 2B and 2C). Each phosphate binds one Sso2081 monomer and forms multiple hydrogen bonds with residues Thr10, Ser11 and Gly104. Of note, Tyr133 from the aforementioned helical insert is also implicated in phosphate binding. These residues are conserved among Sso2081 homologs from different species (Supplementary Fig. 2), suggesting that they may be critical for cA_4_ cleavage.

**Figure 2.**
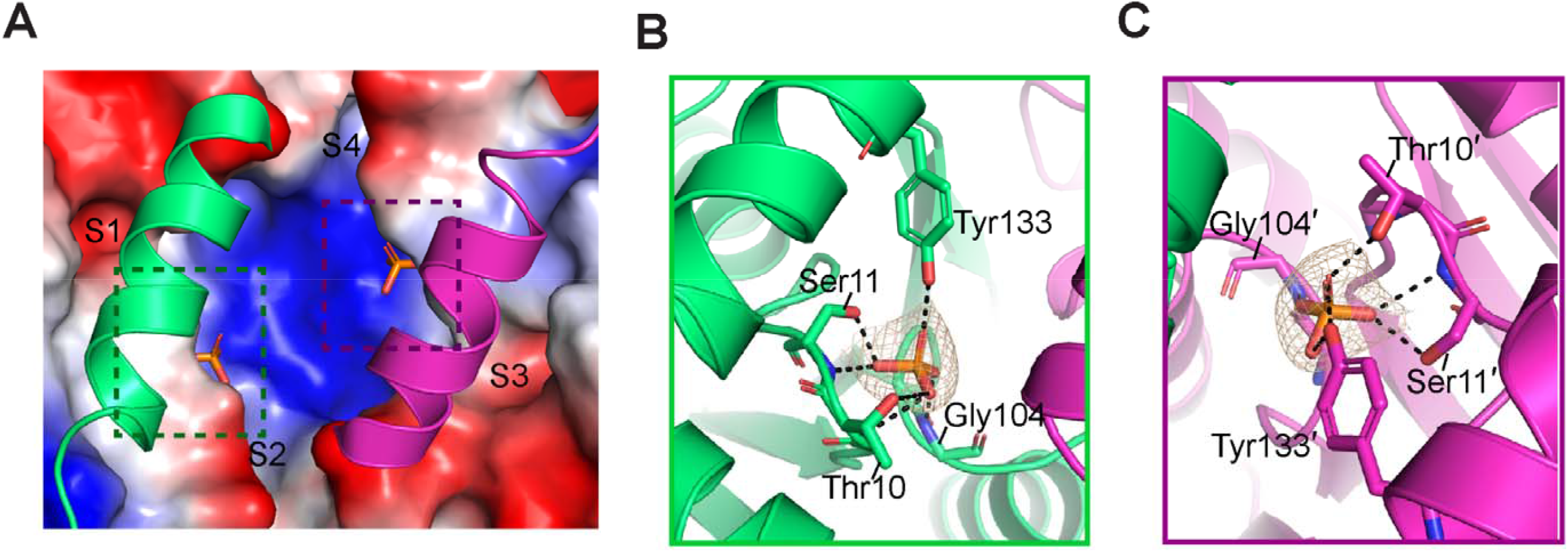
The active sites of Sso2081. **(A)** The surface drawing of the active sites of Sso2081. The helical inserts are shown in cartoon. The surface is colored according to the electrostatic potential (blue, positive; red, negative; white, neutral). The two free phosphates are shown as sticks, and the adenine-binding pockets (S1∼4) are indicated. **(B, C)** The free phosphate-binding sites in monomer-1 (b) and -2 (c), with the 2Fo−Fc electron density map of the bound phosphates shown at 2.0 σ. Key interacting residues are shown as sticks and labeled. Dashed lines indicate hydrogen bonds.

### Structure of the Sso2081:cA_4_ complex in pre-cleavage state

To investigate how Sso2081 recognizes its substrates, we next sought to determine the crystal structure of Sso2081 bound to cA_4_. Initially, we attempted to obtain the structure of Sso2081:cA_4_ complex by soaking the crystals of apo protein into cA_4_-containing solutions but did not succeed. We reasoned that the helical insert hanging over the catalytic pocket might block cA_4_ entry owing to the limited structural flexibility in crystals. With that in mind, we thus pre-incubated Sso2081 with cA_4_ at 4 °C in solution, and crystallized the complex by co-crystallization. The crystals diffracted to a resolution of 3.11 Å, and the structure was determined by molecular replacement using the apo structure as template (Figure 3A). The 2Fo-Fc electron density map clearly showed the presence of cA_4_ in the catalytic pocket (Figure 3B). It is completely enclosed by Sso2081 dimer and adopts a two-fold symmetric architecture. The four phosphodiester bonds do not appear to be cleaved as evidenced by the unambiguous electron density contoured 1.0 σ (Figure 3B), indicating that the structure was determined in the pre-cleavage state. The overall structure of cA_4_-bound Sso2081 is virtually identical to that of apo protein, with an RMSD of 0.73 Å (Supplementary Fig. 4a). The major conformational changes occurred in the helical insert. Upon cA_4_ binding, the two helices get more closer to each other, suggesting a substrate-induced fit mechanism (Figure 3C).

**Figure 3.**
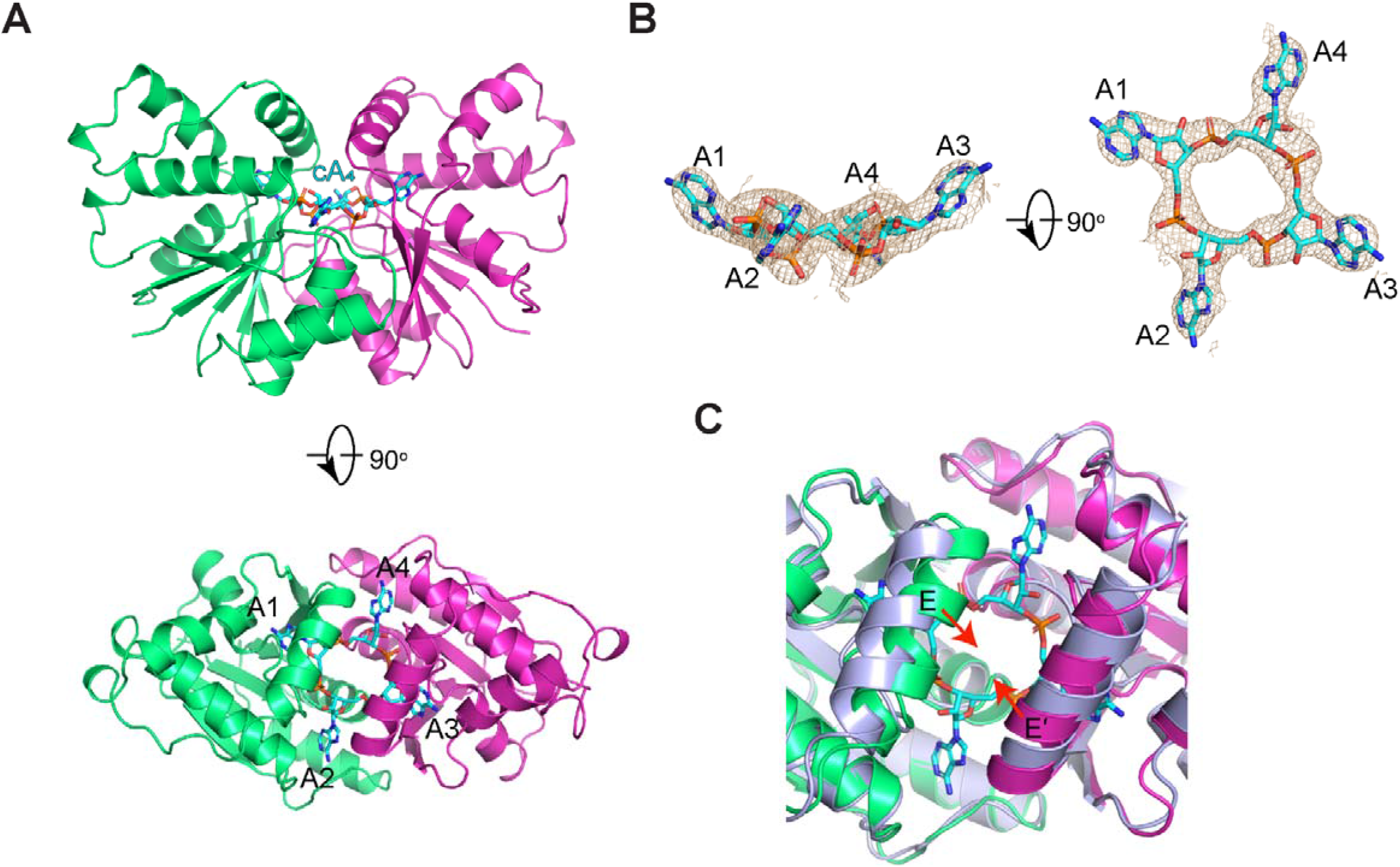
Structure of Sso2081:cA_4_ complex in the pre-cleavage state. **(A)** Cartoon diagram of the crystal structure of Sso2081: cA_4_ complex from side (upper panel) and top (lower panel) views. cA_4_ is shown as cyan stick. **(B)** The 2Fo-Fc electron density map of cA_4_ in the crystal structure, contoured at 1.0 σ. **(C)** Superposition of the structures of apo (light blue) and cA_4_-bound Sso2081. Major conformational changes are indicated by red arrows.

### Substrate recognition

The adenine groups A1 and A3 are respectively bound by monomer-1 and -2 in an acidic cavity under the helical insert, whilst the corresponding groups of A2 and A4 are trapped by two smaller pockets on the dimer interface (Figures 3A and 4A). In the A1 / A3 binding pockets, the adenine base is bound mainly through two hydrogen bonds with the carboxylate side chain of Glu17 (Figure 4B and Supplementary Fig. 4B). Mutation of Glu17 to alanine markedly reduced cA_4_ cleavage (Figures 4E and 4F). On the other hand, the adenine base of A2 / A4 are specifically recognized through the hydrogen-bond network formed by Thr37&, Asp75& and Arg105& (Figure 4C and Supplementary Fig. 4c). Consistent with this, alanine substitution of Asp75 or Arg105 abolished cA_4_ cleavage activity of Sso2081 (Figures 4E and 4F).

**Figure 4.**
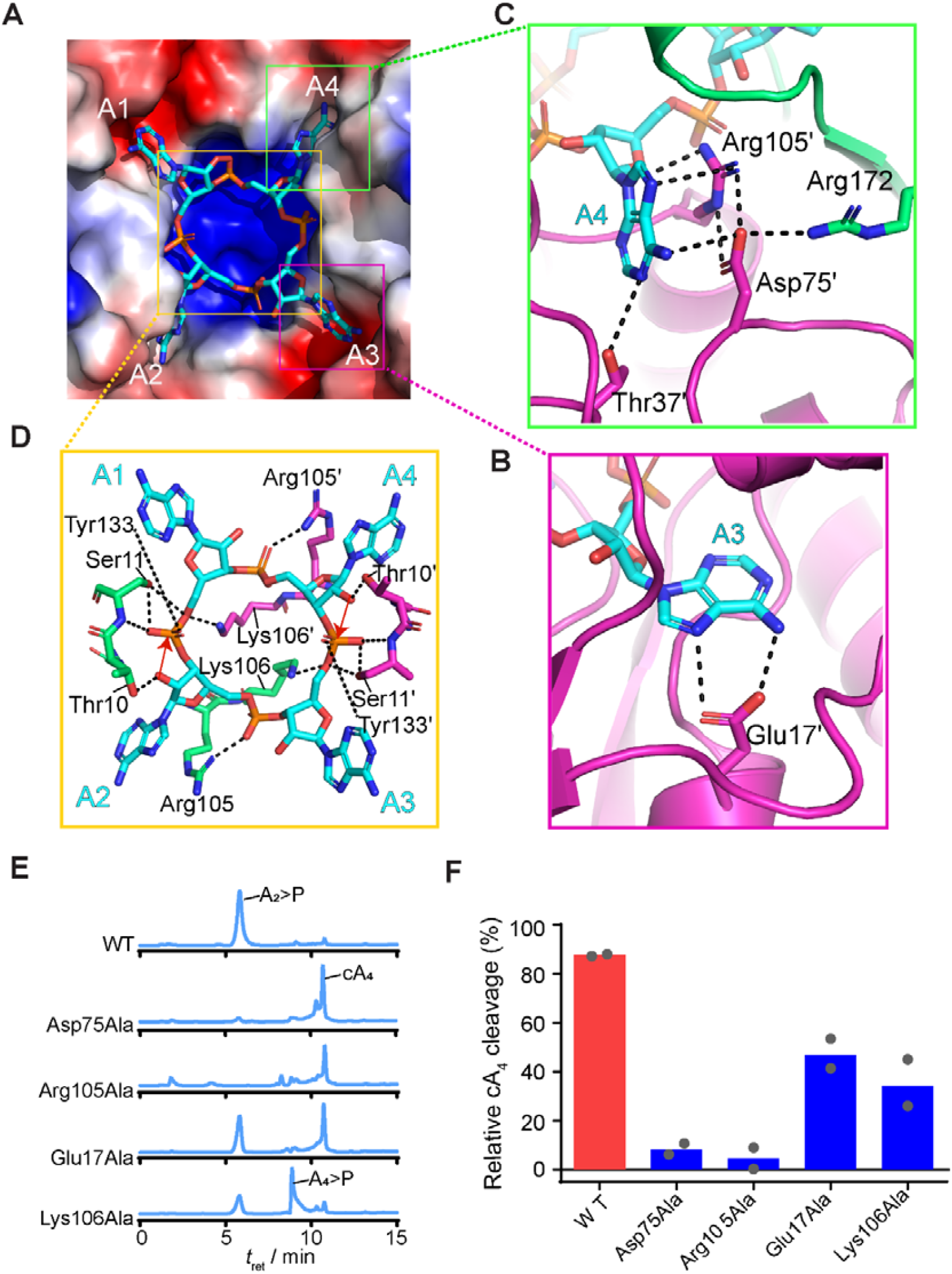
Structural basis of cA_4_ recognition by Sso2081. **(A)** Overall view of cA_4_-binding site on Sso2081 with the helical insert omitted. Sso2081 is shown in surface and cA_4_ is in stick. The surface is colored according to the electrostatic potential, with red, blue and white representing negative, positive and neutral charges, respectively. **(B)** The S3 pockets that recognizes the adenine group of A3. **(C)** The A4 binding site. **(D)** The interactions between Sso2081 and the ribose-phosphate backbone of cA_4_. Dashed lines indicate hydrogen bonds. Residues involved in cA_4_ binding are shown as sticks and labeled. **(E)** LC-MS analyses of cA_4_ cleavage by wild type (WT) Sso2081 or various mutants after incubation at 37 °C for 15 min. The mass spectra of cleavage products by Lys106Ala are shown in Supplementary Fig. 5a. **(F)** Quantitation of cA_4_ cleavage in (E). Values are means of duplicate measurements.

The phosphodiester backbone of cA_4_ docks onto the central basic patch mainly through electrostatic interactions (Figure 4A).In particular, Lys106 on the top of dimeric helices (α4-α4□) from both monomers put their side chains at the center of cA_4_ and point towards the 5□-phosphates of A1 and A3 (Figure 4D). In addition, Arg105 stretches its guanidinium group outside of the tetra-adenylate ring and coordinates the 5□-phosphates of A2 and A4 (Figure 4D). Different to Arg105Ala, the Lys106Ala mutation caused a substantial accumulation of linear intermediates, which was further validated by MS to be 5[-OH-ApApApA-2□, 3□-cyclic phosphate (A_4_>P) (Figure 4E, Supplementary Figs. 1D and 5A). This indicates that Lys106 may be responsible for the transient intermediate stabilization. The 5□-phosphates of A1 and A3 overlap with the two free phosphates in apo Ss2081, and interact with residues Thr10 and Ser11 in a similar way (Figure 4D). Because Ser11 has previously been shown to be catalytically important for cA_4_ cleavage (26), implying that the 5□-phosphates of A1 and A3 are probably the cleaving phosphates.

### Structure of the Sso2081:cA_4_ complex in transient intermediate state

To gain further insights into the catalytic mechanism of cA_4_ cleavage by Sso2081, we next attempted to capture the structure of cleavage intermediate state. Ser11Ala mutation greatly diminished cA_4_ cleavage by Sso2081, resulting in accumulation of A_4_>P intermediate as observed in Lys106Ala mutation (Figure 5A). We therefore pre-incubated the protein of Sso2081 Ser11Ala mutant with cA_4_ at room temperature for over 2 h prior to crystallization. The structure was determined to a resolution of 2.50 Å (Table 1). It is virtually identical to that of the pre-cleavage complex except for the cA_4_ molecule, which was cleaved at the phosphodiester bond between A1 and A2 (Figures 5B and 5C). A structural comparison with the pre-cleavage complex yielded an RMSD of 0.56 Å (Supplementary Fig. 4d), suggesting that cA_4_ cleavage does not induce major conformational changes. Collectively, this structure thereby provides a structural evidence for stepwise cleavage of cA_4_.

**Figure 5.**
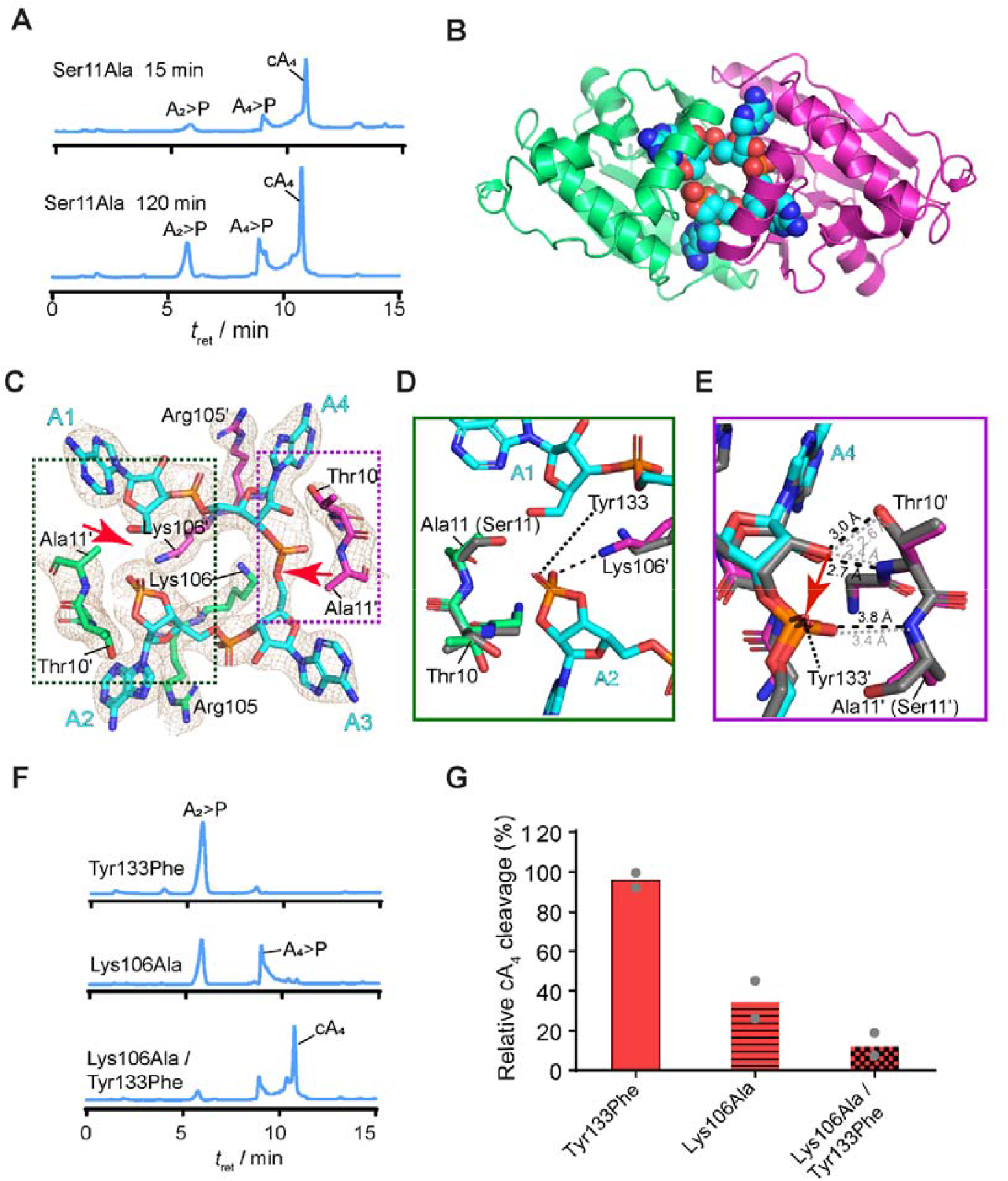
Structure of Sso2081 Ser11Ala:cA_4_ complex in the transient intermediate state. **(A)** LC-MS analysis of cA_4_ cleavage by the Ser11Ala mutant. The mass spectra of cleavage products by Sso2081 Ser11Ala are shown in Supplementary Fig. 5b. **(B)** Top-view of the structure of Sso2081:A_4_>P complex. Sso2081 is shown in cartoon representation and cA_4_ is in sphere. **(C)** The binding site of A_4_>P in Sso2081. The cA_4_ and its interacting residues are shown as sticks and overlaid with the 2Fo-Fc electron density map contoured at 1.0 σ. Red arrows indicate cleavage sites within the cA_4_ ring. **(D, E)** Close-up view of Sso2081 interactions with the cleaved (D) or uncleaved (E) sites of cA_4_, with the superimposition of pre-cleavage state structure (grey). Dashed lines indicate hydrogen bonds. Red arrow indicates nucleophilic attack. **(F)** LC-MS analysis of cA_4_ cleavage by the indicated Sso2081 mutants after incubation at 37 °C for 15 min.. **(G)** Quantitation of cA_4_ cleavage in (f). Values are means of duplicate measurements.

### Mechanism of catalysis

In the structure of transient intermediate state, the cyclic 2□, 3□-phosphate of A2, the product of first cleavage, is stabilized by the side chains of Lys106□ and Tyr133 (Figure 5D). Comparing to the pre-cleavage state, the ε amino group of Lys106□ undergoes a notable conformational change to intimately contact the cleaved phosphate, whereas the corresponding group of Lys106 faces away from the 5□-scissile phosphate of A3 (Figures 5C and 5D). In addition, structure superposition of wild-type Sso2081 with the Ser11Ala mutant demonstrates that Ser11 may involve in the stabilization of 5□-OH of the cleavage intermediate (Figure 5D). On the opposite site where the phosphodiester bond remained uncleaved, the distance between the scissile phosphate of A3 and the amide nitrogen of Ala11 (Ser11) increased from 3.4 Å to 3.8 Å, and the 2□-OH on the ribose also slightly (∼ 0.4 Å) deviated from Thr10 (Figure 5E). These observations thus suggest that Ser11 also plays a role in the alignment of 2□-OH and scissile phosphate for efficient in-line nucleophilic attack.

The helical insert residue Tyr133 from both monomer-1 and -2 coordinate the cleaved and scissile phosphates, respectively. Although single mutation of Tyr133Phe did not result in a noticeable effect, the Tyr133Phe / Lys106Ala double mutation caused a drastic decrease of A_2_>p and A_4_>p products compared to the Lys106Ala mutation (Figures 5F and 5G). This suggests that Tyr133 might play an auxiliary role in the stabilization of either substrate or transient intermediate in the active sites.

### Comparison with the CARF domains of Can2 and Card1

As described above, Sso2081 is structurally most similar to the CARF domains of Can2 and Card1 (Supplementary Fig. 3). Both Can2 (11,21) and Card1 (22) are cA_4_ -activated DNases / RNases, but do not have cA_4_-cleaving activities. Structural comparison showed that the three proteins are quite similar in the architecture of cA_4_ binding sites (Figures 6A-C). The four adenine groups of cA_4_ are bound in a two-fold symmetry, and the phosphodiester backbones are docked onto a glycine loop (highlighted in blue) at the C-terminal end of the central dimeric helices. The cA_4_ centers are occupied by conserved lysines, Lys106 for Sso2081, Lys99 for Can2 and Lys102 for Card1 (Figures 6A-C). In addition, the critical residue equivalent to Arg105 of Sso2081 appears to be substituted by Thr97 of Can2 or Thr101 of Card1 (Figures 6A-C). Despite these similarities, there are remarkable differences in the 5□-phosphate-binding sites of A1 and A3 nucleotides. In Sso2081, the two scissile phosphates are coordinated by the LGTS motif (highlighted in magenta), termed the catalytic loop (Figure 6D). This loop is conserved among ring nucleases from different species including the *S. islandicus* and *S. tokodaii* (Figure 6D). However, the equivalent loops in Can2 and Card1 are different in sequence: LSDH for Can2 and VSEQ for Card1 (Figure 6D). Such sequence diversity in the catalytic loop might lead to the improper alignment of 2□-OH nucleophile and the scissile phosphate in Can2 and Card1, thereby rendering the two proteins inert in cA_4_ cleavage.

**Figure 6.**
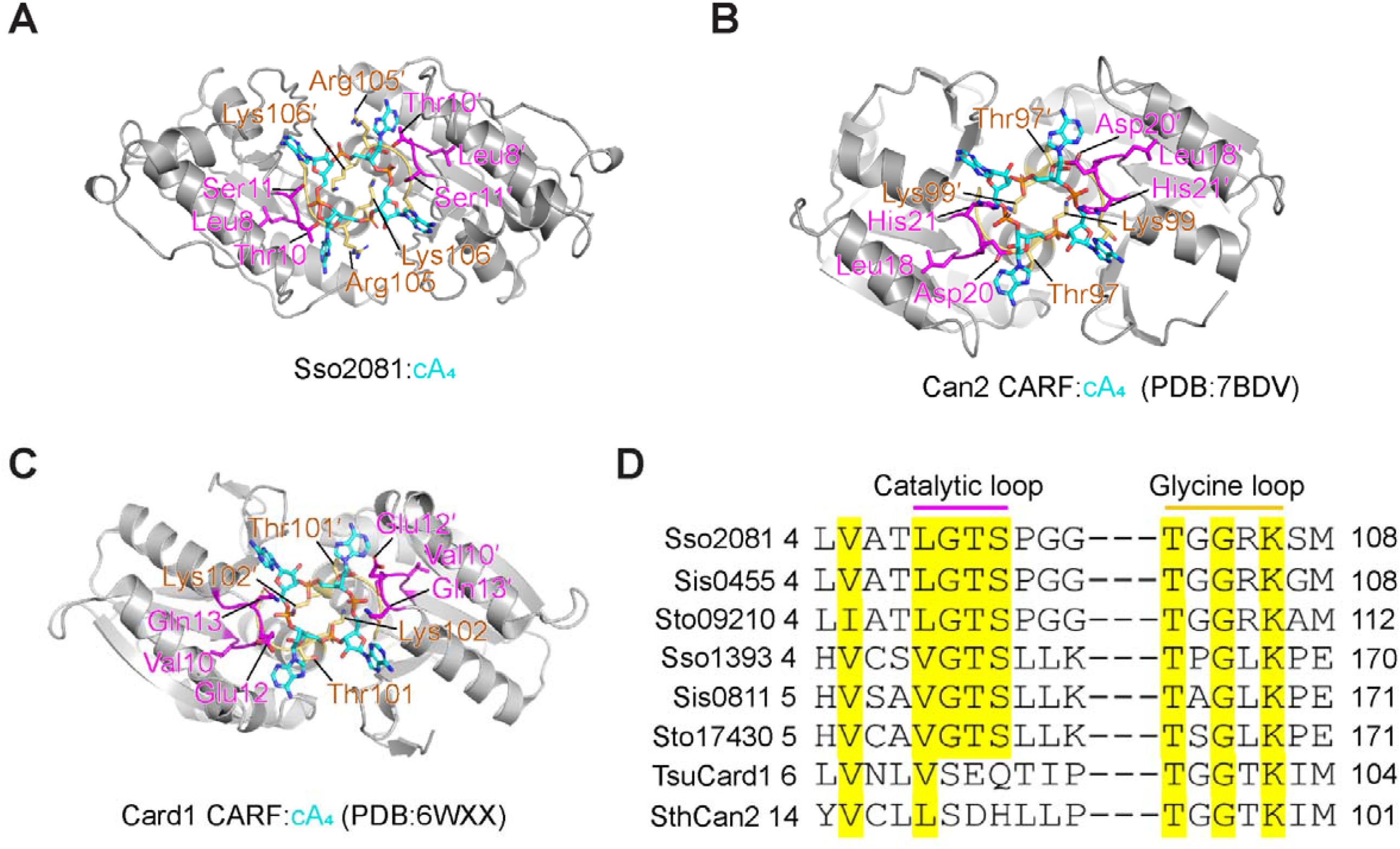
Structural comparisons of Sso2081 with the CARF domains of Can2 and Card1. **(A)** Structure of the Sso2081:cA_4_ complex with the helical insert omitted. **(B)** Structure of Can2 CARF domain in complex with cA_4_ (PDB: 7BDV). **(C)** Structure of Card1 CARF domain in complex with cA_4_ (PDB: 6WXX). **(D)** Sequence alignment between Sso2081 and its structural homologs. The conserve residues are highlighted with yellow shading. The catalytic and glycine loops are marked with magenta and yellow colors, respectively. Sso, *Saccharolobus solfataricus*; Sis, *Sulfolobus islandicus*; Sto, *Sulfolobus tokodaii*; Tsu, *Treponema succinifaciens*;Sth, *Sulfobacillus thermosulfidooxidans*.

## DISCUSSION

The down regulation of cOA signaling in type III CRISPR system is a crucial process to avoid indiscriminate degradation of host RNA. In bacteria and archaea, the cOA degradation is often fulfilled by the standalone ring nucleases, although some Csm6 proteins also exhibit weak cA_4_ or cA_6_ degradation activities (30-32). To date, three subfamilies of standalone ring nucleases have been identified with distinct structural folds (29), the CRISPR-associated ring nuclease 1 (Crn1) (Sso1393 and Ssi0811) (26,27), Crn2 (AcrIII-1) (28) and Crn3 (Csx3) (29). In this study, we describe the crystal structures of Sso2081, alone or bound by cA_4_ in both pre-cleavage and transient intermediate states. We show that Sso2081 is structurally close to the Crn1 subfamily such as Sso1393 (backbone RMSD ∼ 5.4 Å), but is also unique in several aspects (Supplementary Fig. 2). For example, unlike Sso1393, Sso2081 does not contain the C-terminal wHTH domain, which has been shown to auto-inhibit cA_4_ cleavage activity in Sis0811 (homolog of Sso1393) (27). In addition, Sso2081 harbors a unique helical insert which, to our knowledge, has not been observed in other ring nucleases disclosed thus far. The helical insert hangs over the active site cleft and is implicated in transient intermediate stabilization, thereby enhancing cA_4_ cleavage by Sso2081. Previously, Sso2081 has been shown to be 10-fold more active than Sso1393 in cA_4_ cleavage, and hence was suggested to be the major cA_4_-degrading enzyme in the organism of *S. solfataricus* (26). The unique structural properties of Sso2081 revealed herein may, at least in part, explain why it is more active than Sso1393 in cA_4_ cleavage.

Different to many other ring nucleases, the active site of Sso2081 adopts a closed conformation owing to the presence of a helical insert. Upon cA_4_ binding, this helical insert completely encloses the substrate molecule in the catalytic center. Considering the large size of cA_4_ molecule, we propose that the helical insert would undergo a dynamic conformational change to allow substrate entry. Our observation that a major shift occurs in the helical insert upon cA_4_ binding supports this hypothesis.

HPLC-MS analysis show that Sso2081 converts cA_4_ into the final product of A_2_>P. Consistent with this, two free phosphates symmetrically bind the catalytic site of apo Sso2081 and overlap with the two scissile phosphates of cA_4_, supporting a bilaterally symmetrical cleavage. Furthermore, our results provide structural evidences that the two phosphodiester bonds are cleaved in a nick and counter-nick mechanism, as previously proposed for the Csm6 CARF domain (30). Firstly, we detected a substantial accumulation of linear intermediate (A_4_>P) in the reaction products of Ser11Ala and Lys106Ala mutants but not in that of wild type Sso2081, demonstrating that the two sites are cleaved in a stepwise manner and that the two cleavages are not operated independently. Moreover, by introducing a Ser11Ala mutation, we have captured the structure of linear intermediate (A_4_>P) bound state. With this snapshot, we were able to visualize key residues (Lys106 and Tyr133) involved in the stabilization of transient intermediate. We propose that the stabilization of 5□-OH and cyclic 2□, 3□-phosphate at cleaved site would in turn facilitate nucleophile alignment at the opposing site. Because most of the catalytic residues are universally conserved among ring nucleases, the mechanism proposed for Sso2081 might also apply to other members in the family.

The type III CRISPR system employs cOA-activated auxiliary nucleases to eliminate invading DNA and RNA. These nucleases largely contain a cOA-binding CARF domains. While some CARF domain-containing effectors like Csm6 develop self-limiting cOA-degrading activity (30-32), others like the Csx1 (19), Can2 (21) and Card1 (22) only have cOA-binding property. By structural homolog searching on DALI server, we show that Sso2081 is closely related to the CARF domains of Can2 and Card1. For example, they all share a similar architecture in the cA_4_–binding pocket and contain a conserved glycine loop. However, we find that those cA_4_-degrading homologs contain a conserved “VGTS” motif in the 5□-phosphate binding sites of A1 and A3, whereas Can2 and Card1 do not. Combined with the structural and biochemical data, we propose that the VGTS motif, named as catalytic loop, might be a critical determinant of difference in cA_4_ cleavage for CARF domain-containing proteins.

As a summary, in this work, we have determined the crystal structures of Sso2081, alone or bound to cA_4_ in both pre-cleavage and transient intermediate states. These structures together with extensive biochemical analysis provide molecular details of how cA_4_ is specifically recognized and cleaved by Sso2081 in a cooperative nick and counter-nick mechanism. Because of the high sequence and structure conservation, the mechanism proposed herein might also apply to other ring nucleases in the type III CRISPR system. Finally, our structure and sequence analyses provide a new insight to distinguish between cA_4_–degrading and –nondegrading CARF domain-containing proteins.

## ACCESSION NUMBERS

Atomic coordinates and structure factors for the reported crystal structures have been deposited with the Protein Data bank under accession number 7YHL (Apo Sso2081), 7YGH (Sso2081:cA4), 7YGL (Sso2081 S11A:cA4>P).

## ACKNOWLEDGEMENTS

We thank the staffs from BL02U1 and BL19U1 beamlines of National Facility for Protein Science in Shanghai (NFPS) at Shanghai Synchrotron Radiation Facility (SSRF), Shanghai, People’s Republic of China, for assistance with X-ray data collections.

## AUTHOR CONTRIBUTIONS

L. Du performed all the experiments and analyzed the data. Z. Luo performed structure refinement. Z. Lin supervised the project, analyzed the data and wrote the paper. All authors have read and approved the final version of the manuscript.

