## Supplementary material for "Molecular basis of stepwise cyclic tetra-adenylate cleavage by the type III CRISPR ring nuclease Crn1/Sso2081": Supplementary data.docx

**Abstract**

The cyclic oligoadenylates (cOAs) act as second messengers of type III CRISPR immunity system through activating the auxiliary nucleases for indiscriminate RNA degradation. The cOA-degrading nucleases (ring nucleases) provide an ‘off-switch’ regulation of the signaling, thereby preventing cell dormancy or cell death. Here, we describe the crystal structures of the CRISPR-associated ring nuclease 1 (Crn1) from *Saccharolobus solfataricus* (Sso) 2081 in its apo or bound to cA_4_ in both pre-cleavage and transient intermediate states. Sso2081 harbors a unique helical insert that encloses cA_4_ in the central cavity. Two free phosphates symmetrically bind the catalytic site of apo Sso2081 and overlap with the two scissile phosphates of cA_4_, supporting a bilaterally symmetrical cleavage. The structure of transient intermediate state captured by Ser11Ala mutation immediately illustrates a stepwise cleavage of cA_4_ by Sso2081. Our study establishes atomic mechanisms of cA_4_ recognition and degradation by the type III CRISPR ring nuclease Crn1/Sso2081.

**Key words:** Type III CRISPR; cyclic oligoadenylate; cA_4_; ring nuclease; Sso2081.

**Supplementary Figures**

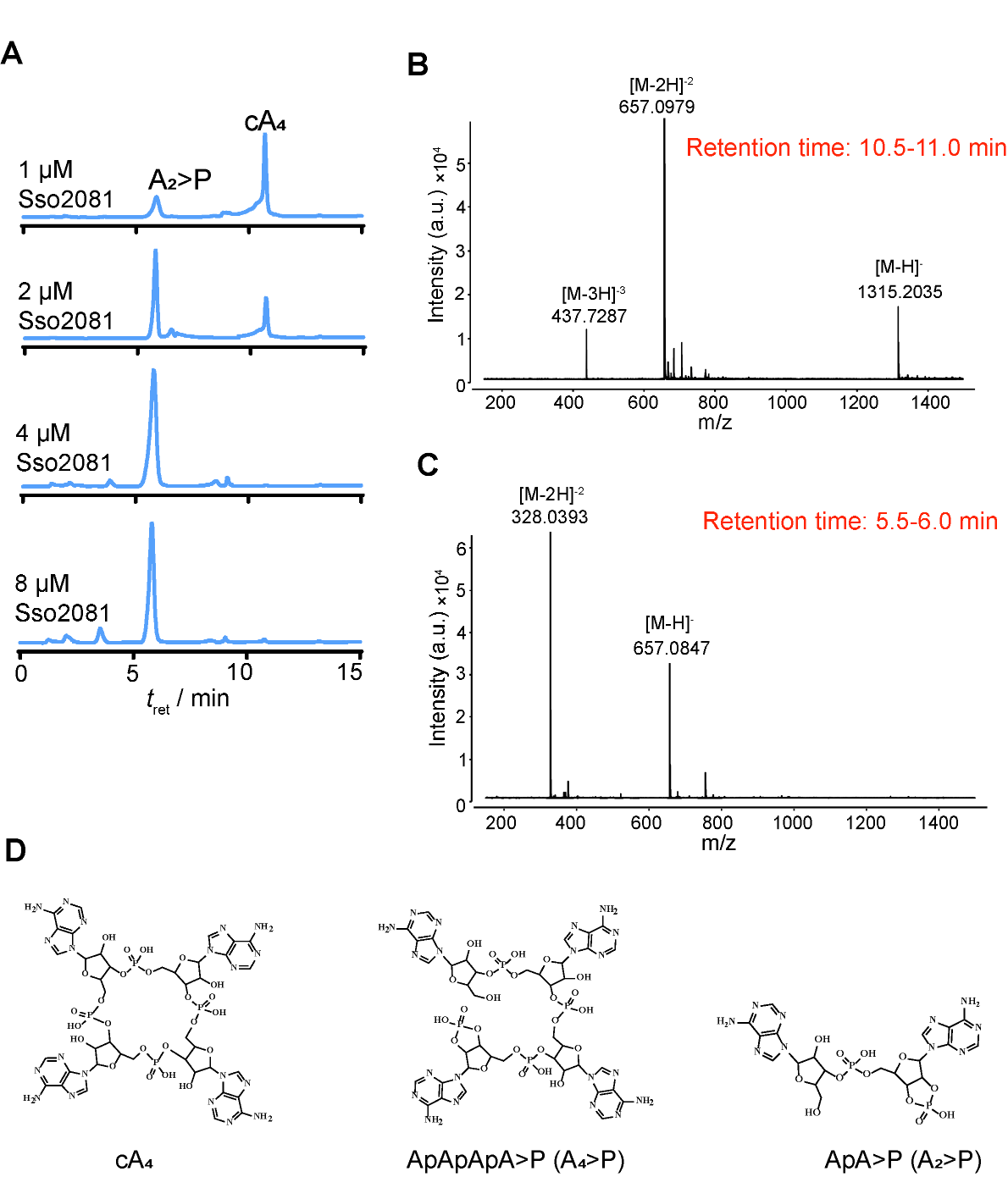

**Supplementary Fig. 1 HPLC and MS analyses of cA_4_ cleavage by Sso2081.** A, HPLC chromatography of cA_4_ cleavage by a series of concentrations of Sso2081. B, C, Mass spectra of the samples eluted from HPLC column at indicated retention time. *m/z* 1315.2035 for cA_4_^-1^; *m/z* 657.0966 for cA_4_^-2^; *m/z* 437.7287 for cA_4_^-3^; *m/z* 657.0847 for A_2_>P^-1^ (ApA>P^-1^); *m/z* 328.0393 for A_2_>P^-2^. D, The chemical structures of cA_4_ and the cleavage products.

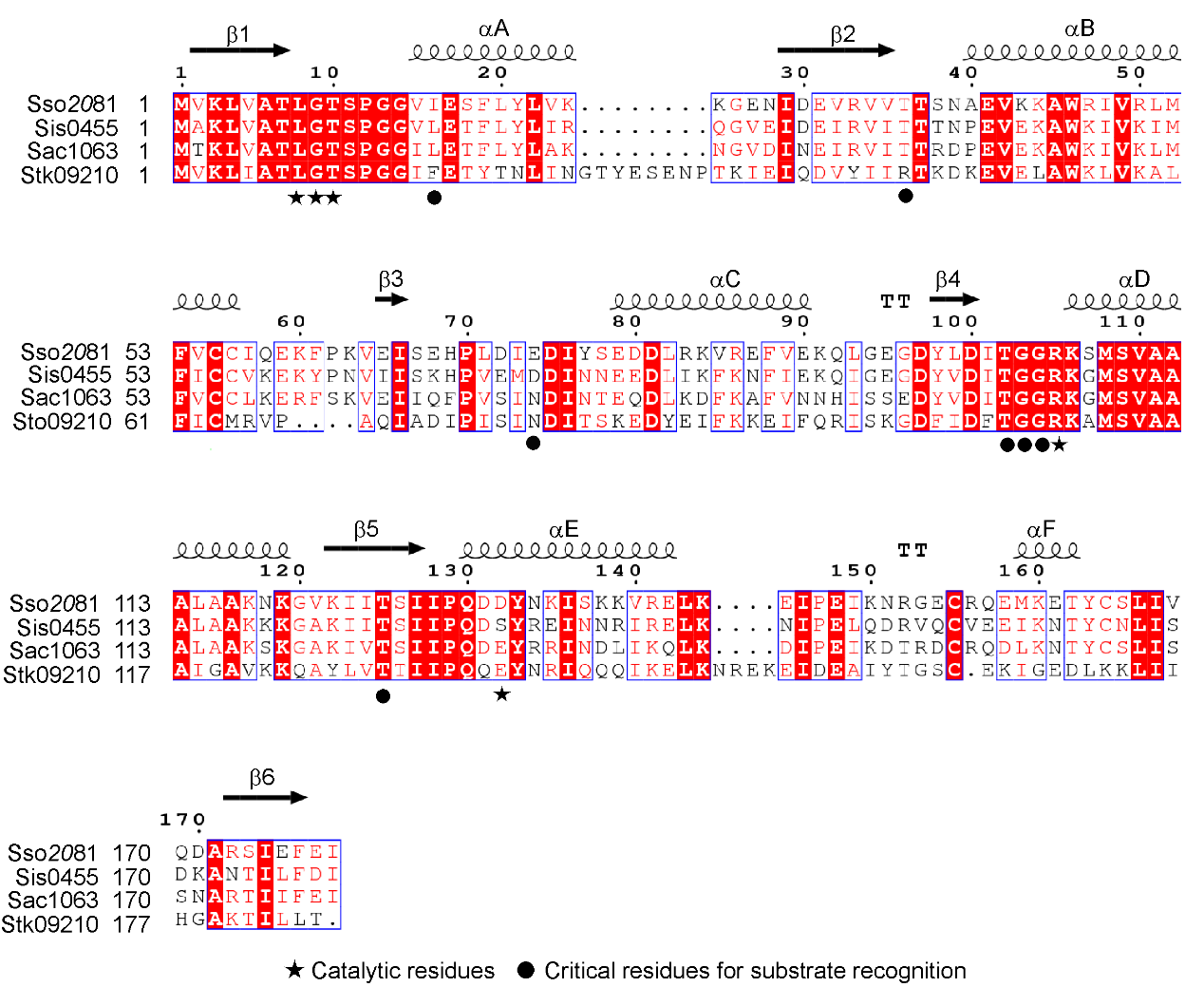

**Supplementary Fig. 2 Sequence alignment of Sso2081 with its structural homologs.** The alignment is generated using the online ESPript 3.0 server. Secondary structural elements of Sso2081 are indicated above the sequences. Abbreviations: Sso, *Saccharolobus solfataricus*; Sis, *Sulfolobus islandicus*; Sac, *Sulfolobus acidocaldarius.* Sto, *Sulfolobus tokodaii*;

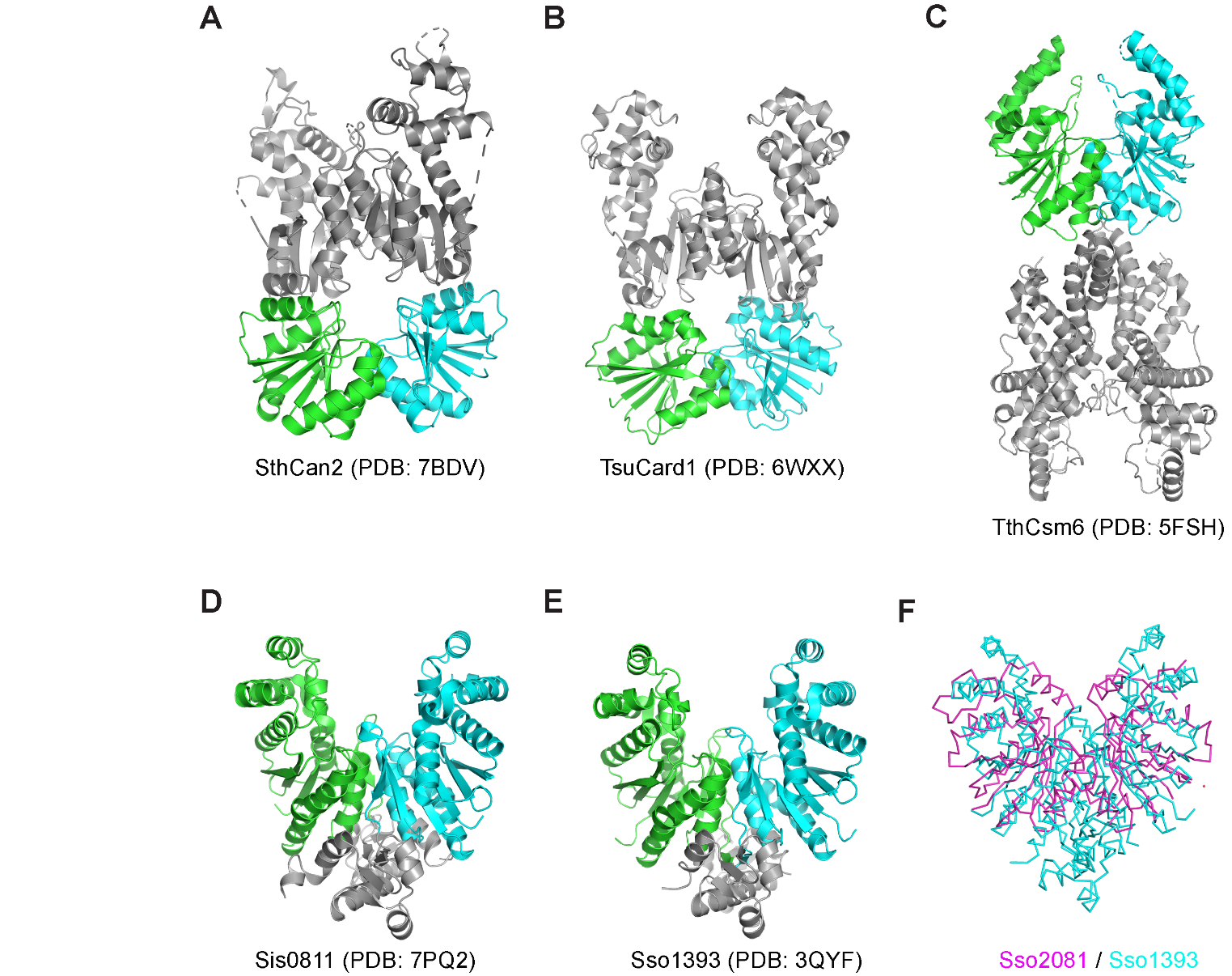

**Supplementary Fig. 3 Comparison of the structures between various CARF domain-containing proteins.** A-E, Structures SthCan2, TsuCard1, TthCsm6, Sis0811 and Sso1393. The structures are shown in cartoon representation. CARF domains are highlighted in green / cyan colors. F, Structural alignment between Sso2081 (magenta ribbon) and Sso1393 (cyan ribbon). Sso, *Sulfolobus solfataricus*; Sth, [*Sulfobacillus thermosulfidooxidans*](https://www.rcsb.org/search?q=rcsb_entity_source_organism.taxonomy_lineage.name:Sulfobacillus%20thermosulfidooxidans); Tsu, [*Treponema succinifaciens*](https://www.rcsb.org/search?request=%7B%22query%22%3A%7B%22type%22%3A%22terminal%22%2C%22service%22%3A%22text%22%2C%22parameters%22%3A%7B%22attribute%22%3A%22rcsb_entity_source_organism.taxonomy_lineage.name%22%2C%22operator%22%3A%22exact_match%22%2C%22value%22%3A%22Treponema%20succinifaciens%20DSM%202489%22%7D%7D%2C%22return_type%22%3A%22entry%22%7D)；Tth, *Thermus thermophilus*.

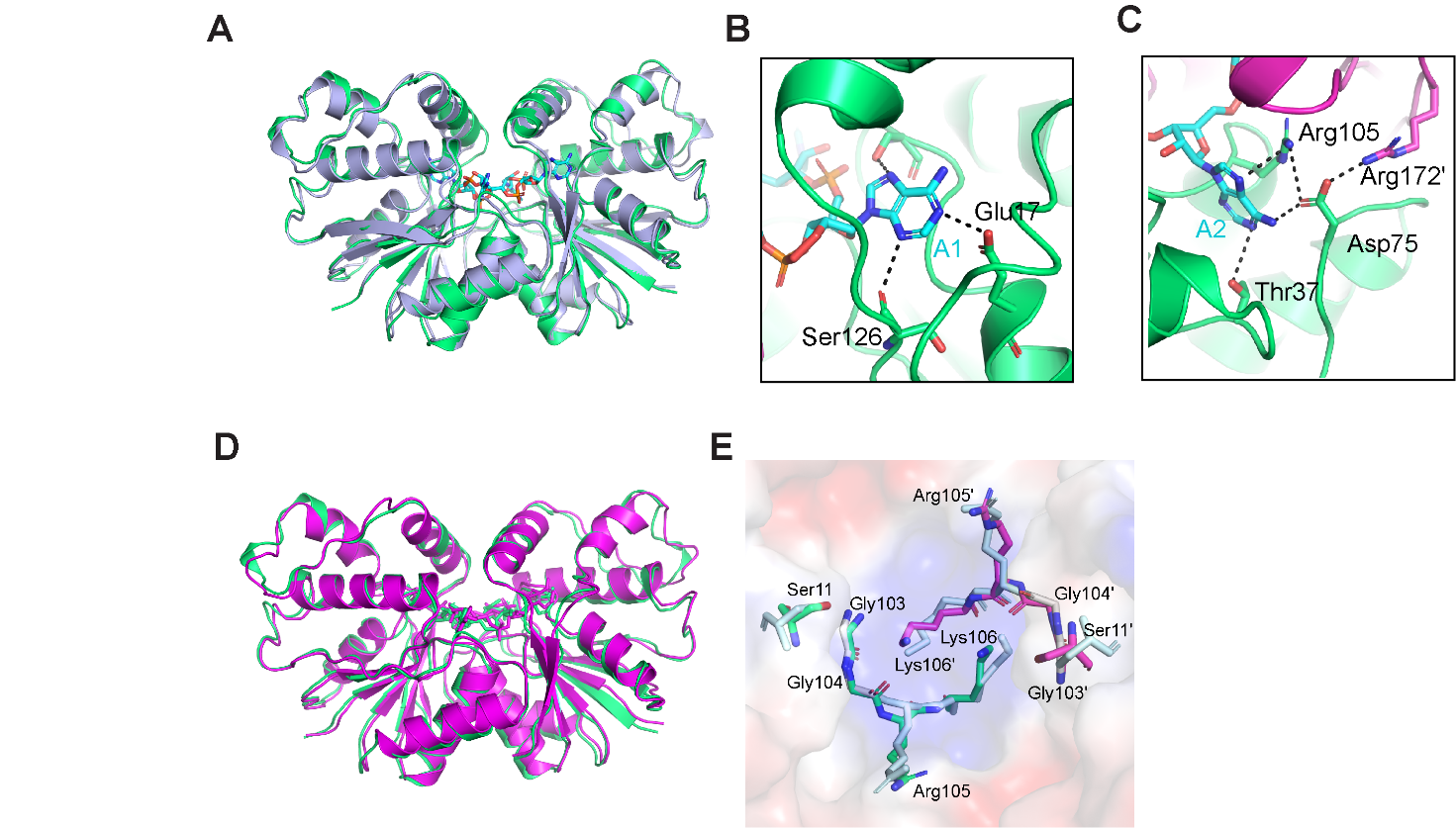

**Supplementary Fig. 4 Structural comparisons of Sso2081 in different states and cA_4_-binding sites.** A, Superposition of the structures of Sso2081 in apo (light blue) and pre-cleavage (green) states. B, C, The binding sites of A1 and A2 groups of cA_4_. D, Superposition of the structures of Sso2081 in pre-cleavage (green) and transient intermediate (magenta) states.

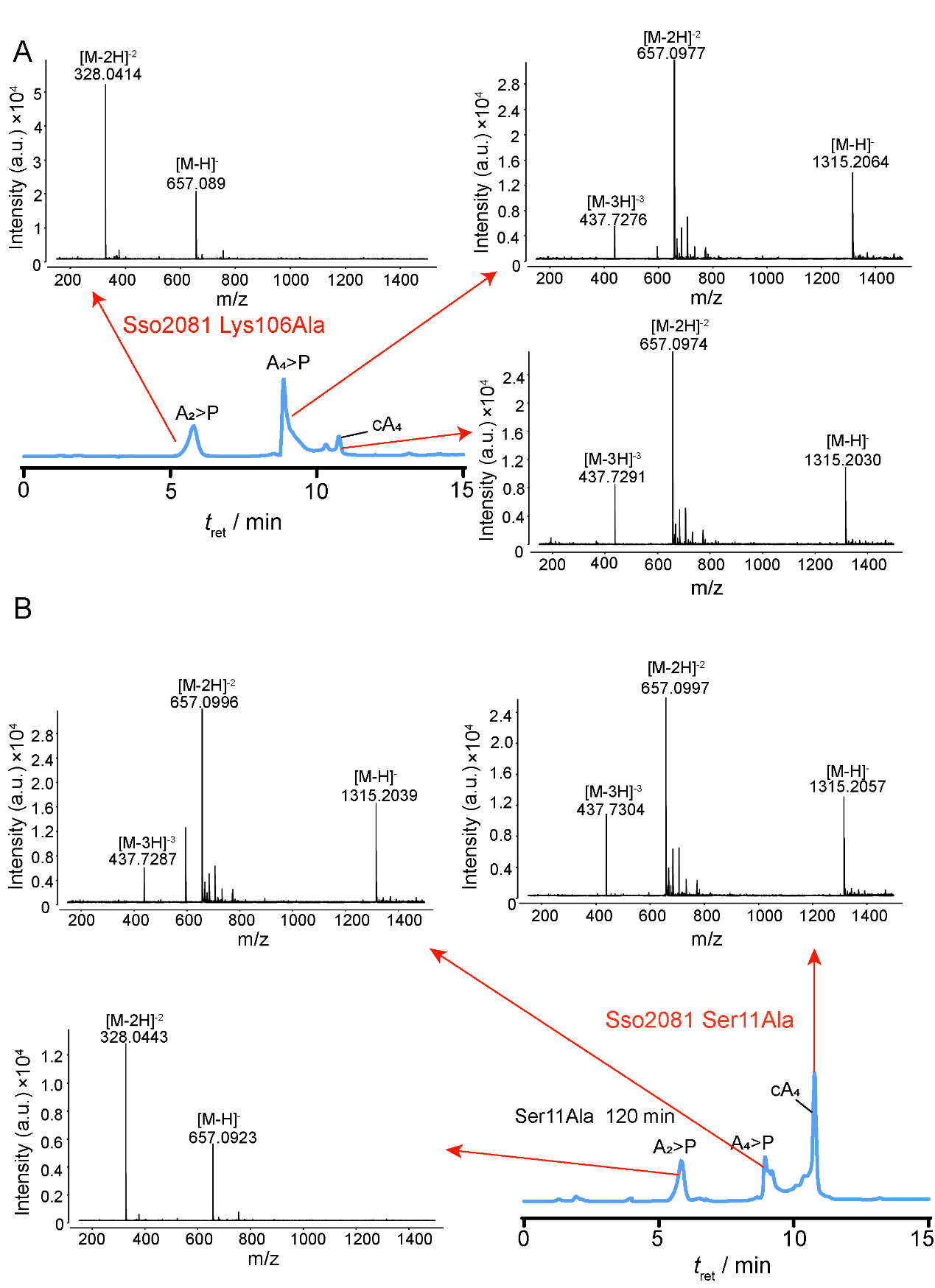

**Supplementary Fig. 5 HPLC chromatography and MS spectra of the cleavage products by the Sso2081 mutants.** A, Sso2081 Lys106Ala, retention time 5.5-6.0: *m/z* 657.0894 for A_2_>P^-1^ (ApA>P^-1^); *m/z* 328.0414 for A_2_>P^-2^; Retention time 9.0-9.5 min: *m/z* 1315.2064 for A_4_>P^-1^ (ApApApA>P^-1^); 657.0977 for A_4_>P^-2^; 437.7276 for A_4_>P^-3^; Retention time 10.5-11.0 min: *m/z* 1315.2030 for cA_4_^-1^; *m/z* 657.0974 for cA_4_^-2^; *m/z* 437.7291 for cA_4_^-3^. B, Sso2081 Ser11Ala, retention time 5.5-6.0: *m/z* 657.0923 for A_2_>P^-1^ (ApA>P^-1^); *m/z* 328.0443 for A_2_>P^-2^; Retention time 9.0-9.5 min: *m/z* 1315.2039 for A_4_>P^-1^ (ApApApA>P^-1^); 657.0996 for A_4_>P^-2^; 437.7287 for A_4_>P^-3^; Retention time 10.5-11.0 min: *m/z* 1315.2057 for cA_4_^-1^; *m/z* 657.0977 for cA_4_^-2^; *m/z* 437.7304 for cA_4_^-3^;
